# High throughput Characterization of *KCNB1* variants Associated with Developmental and Epileptic Encephalopathy

**DOI:** 10.1101/637041

**Authors:** Seok Kyu Kang, Carlos G. Vanoye, Sunita N. Misra, Dennis M. Echevarria, Jeffrey D. Calhoun, John B. O’Connor, Katarina L. Fabre, Dianalee McKnight, Laurie Demmer, Paula Goldenberg, Lauren E. Grote, Isabelle Thiffault, Carol Saunders, Kevin A. Strauss, Ali Torkamani, Jasper van der Smagt, Koen van Gassen, Robert P. Carson, Jullianne Diaz, Eyby Leon, Joseph E. Jacher, Mark C. Hannibal, Jessica Litwin, Neil R. Friedman, Allison Schreiber, Bryan Lynch, Annapurna Poduri, Eric D. Marsh, Ethan M. Goldberg, John J. Millichap, Alfred L. George, Jennifer A. Kearney

**Author notes:** Denotes equal contribution (Seok Kyu Kang, Carlos G. Vanoye). **Corresponding Author**: Jennifer A. Kearney, Department of Pharmacology, 320 East Superior St, Searle 8-510, Northwestern University, Chicago, IL 60611.

## Abstract

Pathogenic variants in *KCNB1*, encoding the voltage-gated potassium channel Kv2.1, are associated with developmental and epileptic encephalopathies (DEE). Previous functional studies on a limited number of *KCNB1* variants indicated a range of molecular mechanisms by which variants affect channel function, including loss of voltage sensitivity, loss of ion selectivity, and reduced cell-surface expression. We evaluated a series of 17 *KCNB1* variants associated with DEE or neurodevelopmental disorder (NDD) to rapidly ascertain channel dysfunction using high-throughput functional assays. Specifically, we investigated the biophysical properties and cell-surface expression of variant Kv2.1 channels expressed in heterologous cells using high-throughput automated electrophysiology and immunocytochemistry-flow cytometry. Pathogenic variants exhibited diverse functional defects, including altered current density and shifts in the voltage-dependence of activation and/or inactivation, as homotetramers or when co-expressed with wild-type Kv2.1. Quantification of protein expression also identified variants with reduced total Kv2.1 expression or deficient cell-surface expression.

Our study establishes a platform for rapid screening of functional defects of *KCNB1* variants associated with DEE and other NDDs, which will aid in establishing *KCNB1* variant pathogenicity and may enable discovery of targeted strategies for therapeutic intervention based on molecular phenotype.

## INTRODUCTION

*De novo* variants in a diverse range of genes constitute a significant cause of developmental and epileptic encephalopathies (DEE) and other neurodevelopmental disorders (NDD)^1,2^. Despite an increasing number of genes identified for DEE and increased implementation of clinical genetic testing, many variants remain variants of uncertain significance (VUS) that are difficult to interpret and/or act on. One powerful approach to bridge knowledge gap between the genetics and the molecular pathology is to obtain functional data^3^, which can then provide additional evidence for interpretation of variant pathogenicity and may offer information (e.g, loss-of-function vs. gain-of-function) to guide treatment strategies. However, due to the low throughput of many standard functional assays, functional annotation of variants has become a major bottleneck in the field. The development of high-throughput functional assays is a necessary step for the experimental validation of the large volume of novel variants being identified due to increasing clinical genetic testing.

*KCNB1* encodes the Kv2.1 voltage-gated potassium (K^+^) channel α-subunit that conducts delayed rectifier K^+^ currents^4^, a key modulator of membrane repolarization in electrically excitable cells, including neurons of various subtypes^5^. *Kcnb1*-deficient mice display enhanced seizure susceptibility and behavioral hyperexcitability, indicating that Kv2.1 acts as a homeostatic suppressor of heightened neuronal activity^6^. *De novo* missense variants of *KCNB1* are associated with DEE^7–13^, which includes a wide spectrum of clinical phenotypes including multiple seizure types, developmental delay (DD), and neuropsychiatric sequelae. Correlational analyses based solely on the substituted amino acid type and position have proven insufficient for genotype-phenotype association^11^, highlighting the need for experimental studies to determine functional effects of additional variants. To date, seven *KCNB1* variants associated with DEE, all located in the pore (S347R, T374I, V378A, G379R, G401R) and voltage-sensor (I199F, R306C) domains, have been shown to exhibit altered potassium current density, voltage-dependence, and/or ion selectivity^7–9,14^. However, determining the functional consequences of a larger series of variants is necessary to define the full range of dysfunction and to broaden the range of associated clinical phenotypes studied.

In this study, we performed high-throughput functional studies of 19 *KCNB1* variants (17 missense, 1 frameshift, 1 nonsense) in order to assess their effect on protein function and provide functional evidence for weighing potential pathogenicity. Several broad categories of channel dysfunction (i.e. altered peak current density, voltage-dependence, protein expression) were identified, and most of the examined variants confer some degree of loss-of-function. Our results provide insight into functional pathogenicity for a large series of *KCNB1* variants. This will aid in prioritization for development of more complex experimental models (i.e. knock-in mouse or iPSC) by identifying a subset of representative variants based on underlying pathophysiologic mechanisms. Uncovering a range of *KCNB1* functional defects will help define genotype-phenotype relationships by adding a molecular phenotype to the genotypes and may ultimately enable development of targeted treatment strategies for individuals with *KCNB1* DEE.

## MATERIALS AND METHODS

### Study Subjects

Clinical and genetic information was retrospectively collected for individuals with DEE or global DD of unknown cause and a potentially pathogenic *KCNB1* variant identified by clinical genetic testing as part of routine care. Clinical and genetic information was collected under approval of the Ann & Robert H. Lurie Children’s Hospital of Chicago Institutional Review Board (IRB), the IRB of collaborating institutions, or through collection of de-identified medical information deemed non-human subjects research by the Northwestern University Institutional Review Board (IRB). Consent for release of deidentified information was obtained from the parents/guardians in accordance with local institutional policies. In addition, 11 *KCNB1* variants were obtained from the literature^8–13,15,16^ and variant databases^17–19^. Functional studies had not been previously reported for 17 of the 19 variants described in this study.

### Cell Culture

Chinese hamster ovary cells (CHO-K1, CRL 9618, American Type Culture Collection, Manassas VA, USA) were grown in F-12K nutrient mixture medium (GIBCO/Invitrogen, San Diego, CA, USA) supplemented with 10% fetal bovine serum (FBS, ATLANTA Biologicals, Norcross, GA, USA), penicillin (50 units·mL^−1^), streptomycin (50 μg·mL^−1^) at 37°C in 5% CO_2_. Unless otherwise stated, all tissue culture media were obtained from Life Technologies, Inc. (Grand Island, NY, USA).

### Plasmids

Full-length human wild-type KCNB1 (Kv2.1) cDNA engineered in the plasmid pIRES2-DsRed-MST was previously described^7,14^. Mutations were introduced by QuikChange mutagenesis (Agilent, Santa Clara, CA, USA) using pIRES2-DsRed-MST-KCNB1-WT as template. All recombinant cDNAs were sequenced in their entirety to confirm the presence of the desired modifications and the absence of unwanted mutations in the cDNA. Endotoxin-free plasmid DNA from sequence-verified clones was isolated using the Nucleobond Xtra Midi EF Kit (Macherey-Nagel, Düren Germany), ammonium acetate precipitated and re-suspended in MaxCyte buffer (MaxCyte Inc., Gaithersburg, MD, USA). For cotransfection experiments, pIRES2-DsRed-MST-KCNB1-WT was engineered to replace DsRed-MST with smGFP using Gibson assembly to generate pIRES2-smGFP-KCNB1-WT. For cell-surface labeling of Kv2.1 proteins, a 2x tandem HA tag (YPYDVPDYAYPYDVPDYA) was inserted between phenylalanine 220 and glycine 221, followed by removal of downstream of IRES element and the reporter fluorophore, resulting in plasmid pCMV-KCNB1-HA-WT.

### Heterologous Expression

Transient expression of wild-type (WT) and variant *KCNB1* in CHO-K1 cells was achieved by electroporation using the Maxcyte STX (MaxCyte Inc.) system as per manufacturer instructions. Briefly, cells grown to 70-80% confluence were harvested using 0.25% trypsin-EDTA, washed once with electroporation buffer (EBR100; MaxCyte Inc.) and then re-suspended in the same buffer to a density of 100 × 10^6^ viable cells/mL. For each electroporation, *KCNB1* cDNA (7.5-10μg) was added to 100 μL of cell suspension and the mixture was transferred to an OC-100 (OC-100R10; MaxCyte Inc.) processing assembly, then electroporated using the preset CHO or CHO-PE (protein-expression) protocol. Immediately after electroporation, 10 μL of DNaseI was added to DNA-cell suspension and the whole mixture was then transferred into a 60mm dish and incubated for 30 min at 37°C in 5% CO_2_. Following incubation, the cell suspension was gently re-suspended in culture media, transferred to a T75 tissue culture flask, and grown for 36 hours at 37°C in 5% CO_2_. After this period, cells were harvested and then frozen in liquid N_2_. Co-expression of WT and variant was achieved by co-electroporation of pIRES2-DsRed-MST-KCNB1-WT or -variant (5 μg) plus pIRES2-EGFP-KCNB1-WT (5 μg) using the same protocol.

### Cell Preparation for Automated Patch Clamp Recording

Cells were thawed and incubated for 24 hours at 37°C in 5% CO_2_. Prior to the experiment, cells were harvested using 0.25% trypsin-EDTA in cell culture media. Cell aliquots were used to determine cell number and viability by automated cell counting (Vi-Cell, Beckman Coulter, Indianapolis, IN, USA). Cells were diluted to a concentration of 200,000 total cells/mL with external solution (see below for composition) and then allowed to recover for 40 minutes at 15°C while shaking on a rotating platform at 200 rpm.

### Automated Patch Clamp Recording

Automated patch clamp recording was performed using the SyncroPatch 768PE system (Nanion Technologies, Munich, Germany). The external solution contained (in mM): 140 NaCl, 4 KCl, 1 MgCl_2_, 2 CaCl_2_, 5 glucose, and 10 HEPES, pH 7.4. The internal solution contained (in mM): 60 KF, 50 KCl, 10 NaCl, 20 EGTA, 10 HEPES, pH 7.2. Pulse generation and data collection were done with PatchController 384 V1.3.0 and DataController384 V1.2.1 software (Nanion Technologies) and wholecell currents were acquired at 5 kHz and filtered at 1 kHz. Currents were not leak subtracted. Access resistance and apparent membrane capacitance were estimated using built-in protocols in the PatchController software. Whole-cell currents were measured at room temperature from a holding potential of −80 mV and elicited with depolarizing steps (500 ms) from −100 to +60 mV (10 mV steps) followed by a 250 ms step to −30 mV or 0 mV (tail currents). Background currents were removed by digital subtraction of whole-cell currents recorded from non-transfected CHO-K1 cells off-line. Current amplitudes were analyzed from all cells with seal resistance ≥0.5Gω, series resistance ≤20Mω and cell capacitance >2pF. Electrically unstable cells with loss of seal or voltage control during experiments were excluded from the analysis. The peak current was normalized for cell capacitance and plotted against voltage to generate peak current density–voltage relationships. Voltage-dependence of activation was determined by plotting tail currents normalized to the largest tail current amplitude against the depolarizing test potential (−100 to +60 mV). The normalized G-V curves were fit with the Boltzmann function: G = 1 / (1 + exp[(V-V_½_) / k] to determine the voltage for half-maximal channel activation (V_½_) and slope factor (k). The voltage-dependence of inactivation was assessed following a 5 s pre-pulse from −100 to +40 mV (10 mV steps) followed by a 250 ms step to +60 mV. The normalized peak currents measured at the +60 mV test potential were plotted against the pre-pulse voltage and fit with the Boltzmann function (I/Imax = 1 / (1 + exp[(V-V_½_) / k]) to determine the voltage for half-maximal channel inactivation (V_½_) and slope factor (k). Voltage-dependence of channel activation or inactivation were determined only from cells with positive outward current values following background subtraction. Statistical outliers (values that fell outside the group mean ± 3 standard deviations range) were excluded from the final results.

### Immunocytochemistry-Flow Cytometry

Cells were electroporated as described above and incubated for 24 hours at 37°C in 5% CO_2_ on 60mm tissue-culture dishes. For analysis of total and cell surface expression, cells were washed twice with PBS (+Mg^2^+/Ca^2^+) and incubated with αHA-Alexa488 (16B12, Thermo Fisher, Waltham, MA, USA; 1:250 dilution in FACS buffer (1% FBS, 0.05% NaN3 in PBS, pH 7.4)) for 1h at room temperature. Cells were then harvested with Accutase, collected by centrifugation at 1000g for 2 min, and then were subsequently kept on ice and protected from light for the duration of the experiment. Cells were washed twice with FACS buffer and fixed/permeabilized using Fix&Perm buffer (BD Biosciences, San Jose, CA, USA) according to manufacturer’s instructions. Briefly, cells were washed twice with Perm buffer (BD sciences) for 3 min and incubated with Alexa647-conjugated anti-Kv2.1 mouse monoclonal antibody (K89/34, Neuromab, Antibodies Inc., Davis, CA; A20186 Thermo Fisher; 1:200 dilution in Perm buffer with 3% NGS) for 1h. Cells were washed 4 times with Perm buffer followed by 2 times with FACS buffer. Fluorescence signals were then measured using a benchtop flow cytometer (CytoFlex, Beckman Coulter) and analyzed with CytExpert software (Beckman Coulter) and FlowJo software (FlowJo LLC). Non-transfected CHO cells prepared in the same way served as negative controls. Light-scatter based gates (FSC/SSC) were used to exclude non-viable cells and cell debris. A total of 10,000 events were analyzed for each Kv2.1 variant per experiment.

For independent analyses of Kv2.1 total protein expression (Fig. 3C&E), frozen aliquots of cells electroporated as described above were thawed and cultured for 24 hours, and then directly harvested with Accutase, without the cell-surface protein labeling steps. Percent-positive variant-expressing cells (excited by 638nm laser) were normalized to WT-expressing cells tested on the same day. Histograms were aligned to correct for between-run variability in fluorescence intensity; y-axes were uniformly set at a maximum of 250 and 350 event counts (for Fig. C and E respectively) across conditions.

### Cell Surface Biotinylation and Immunoblotting

Frozen aliquots of CHO-K1 cells electroporated with homotetramer WT or variant KCNB1 channels were thawed and grown under the same conditions used for automated patch clamp recording. Cell surface proteins were labeled with cell membrane impermeable Sulfo-NHS-Biotin (#21217, Thermo Fisher, Waltham, MA), quenched with 100 mM glycine, lysed with RIPA lysis buffer (#89901, Thermo Fisher) containing complete protease inhibitors (04693124001, Roche, Indianapolis, IN, USA) and clarified by centrifugation. Supernatant samples were collected and an aliquot was retained as the total protein fraction. Biotinylated surface proteins were recovered by incubating 200 μg of total protein with streptavidin-agarose beads (20347, Thermo Fisher) and eluting in Laemmli sample buffer. Total (5 μg per lane) and surface fractions were analyzed by immunoblotting using mouse anti-Kv2.1 (1:250; Neuromab ASC-003), mouse anti-transferrin receptor (1:500; clone H68.4; #13-6800, Thermo Fisher), and rabbit anti-calnexin (1:250; H70, sc-11397; Santa Cruz Biotechnology, Santa Cruz, CA, USA) primary antibodies. Alexa Fluor 680 and 790-conjugated goat anti-rabbit immunoglobulin G (IgG; 1:20,000, 111-625-144, Jackson ImmunoResearch, West Grove, PA, USA) and goat anti-mouse IgG (1:20,000, 115-655-146, Jackson ImmunoResearch) secondary antibodies were used.

Blots were probed initially for transferrin receptor and calnexin in multiplex, followed by stripping with Restore Western Blot Stripping Buffer (21059, Thermo Fisher) and then reprobing for Kv2.1. To control for protein loading, Kv2.1 bands were normalized to the corresponding transferrin receptor band. Blots were imaged using the Odyssey CLx system (LI-COR, Lincoln, NE, USA) and densitometry was performed using ImageStudio software (LI-COR). Normalized total, surface, and surface/total protein ratio results were derived from 3 independent experiments. Calnexin immunoreactivity was present in total protein lysates and absent from the cell surface fraction, confirming the selectivity of biotin labeling for cell surface protein.

### Data Analysis

Data were analyzed and plotted using a combination of DataController384 V1.2.1 (Nanion Technologies), Microsoft Excel, SigmaPlot 2000 (Systat Software, Inc., San Jose, CA, USA), GraphPad Prism (GraphPad Software, San Diego, CA, USA) and OriginPro 2018 (OriginLab, Northampton, MA, USA). Statistical differences between groups were evaluated by One-way ANOVA followed by Dunnett’s post-hoc analyses (for groups >2) and unpaired t-tests (for two groups) for comparisons against control (WT) using GraphPad Prism7. P-values are listed in the text, figure legends or tables. Normalizations as well as statistical comparisons to WT were conducted per plate (for high-throughput electrophysiology) or per experiment (for protein expression analyses) to account for potential batch effects. For voltage-dependence in co-expression experiments for which multiple batches were run, the obtained parameters (V_1/2_ or k) could not be subject to percentile normalization; thus, statistical tests were conducted between average values of WT and each variant using Brown-Forsythe and Welch ANOVA test (unequal variance assumption) with Dunnett’s T3 multiple comparisons. Data fits with Boltzmann’s equation for Table 2 were done for each individual cell using non-linear curve fitting in OriginPro2018; solid fit lines presented in Fig. 4 represent fits of averaged data. Whole-cell currents are normalized for membrane capacitance and results are expressed as mean ± S.E.M. The number of cells used for each experimental condition is given in the Tables or Figure legends.

Principal component analysis (PCA) was performed using ClustVis^20^, with principal components (PC) calculated using the non-linear iterative partial least square (Nipals) algorithm. A total of 6 dimensions from the homomeric condition were compared: Ipeak density, V_1/2_ and *k* for activation and inactivation, cell-surface/total expression ratio. Prior to PCA, data were normalized with respect to the WT by percentile value or Z-score.

### Data availability

The data that support the findings of this study are available from the corresponding author, upon reasonable request.

## RESULTS

### *KCNB1* Variant *In Silico* Analysis and Associated Clinical Phenotypes

*KCNB1* is intolerant of variation, with increased constraint for missense variants and intolerance for loss-of-function variation^18^. Examination of missense single nucleotide variations (SNVs) in the gnomAD database^21,22^ reveals a paucity of missense SNVs within the voltage-sensor and pore domains of the *KCNB1,* whereas all pathogenic variants reported to date are clustered in these regions^7–14^. The altered residues reported in this study all show a high degree of evolutionary conservation (Fig. 1A) and are mostly located in transmembrane domains of the Kv2.1 channel protein (Fig. 1B-C). Serine 202, threonine 210, arginine 306 and arginine 312 are located in the voltage-sensing domain, with arginines 306 and 312 being critical, positively-charged residues in the S4 segment that move in response to voltage changes^23^. Arginine 325 and glutamine 330 are located in the S4-S5 linker, which is critical for electromechanical coupling between the voltage-sensing and pore domains^24^. Tryptophan 370, proline 385, lysine 391, and phenylalanine 416 are located in the pore domain, with valine 378 and glycine 381 located within the selectivity filter^25^. Nonsense variants (frameshift at tyrosine 274 and stop codon at tyrosine 533) are not predicted to undergo nonsense-mediated mRNA decay, as the premature stop codons are located at the last exon, and thus have potential to generate a truncated protein. All disease-associated variants had a scaled CADD score of >23, indicating that they rank among the top 1% of deleterious variants in the human genome (Table 1)^26^.

**Table 1.** Clinical features of individuals with *KCNB1* missense variants. Heterozygous de novo variants were reported as pathogenic or likely pathogenic, with the exception of a single variant determined to be non-pathogenic following parental testing (grey shading).

| Case No. | Category | Sex | Developmental Stage at Seizure Onset (Age) | Seizure Types | Developmental Phenotype | Variant (hg19; NM_004975) | Prediction<br>SIFT<br>PolyPhen<br>CADD Score | gnomAD MAF | Ref |
| --- | --- | --- | --- | --- | --- | --- | --- | --- | --- |
| 1 | DEE | Male | Toddler (2 yo) | Complex febrile; subclinical seizures during sleep | global DD; language impairment; ASD | chr20:47991492 G/A p.S202F | Deleterious<br>Probably damaging<br>29.4 | not present | 11 |
| 2 | DEE | Male | Infant (4 mo) | Myoclonic, atonic, GTC, focal dyscognitive with motor signs, subclinical seizures during sleep; history of ESES | global DD | chr20:47991468 G/T p.T210K | Deleterious<br>Probably damaging<br>28.1 | not present | 11 |
| 3 | DEE | Female | Childhood (7 yo) | GTC | global DD; ID | chr20:47991468 G/A p.T210M | Deleterious<br>Probably damaging<br>28.2 | not present | 11 |
| 4 | DEE | Male | Infant (7 mo) | Atypical absence, myoclonic, GTC | global DD; moderate ID | chr20:47991468 G/A p.T210M | Deleterious<br>Probably damaging<br>28.2 | not present | This report |
| 5 | DEE | Female | Infant (12 mo) | Atypical absence, GTC | global DD; severe ID | chr20:47991468 G/A p.T210M | Deleterious<br>Probably damaging<br>28.2 | not present | 16 |
| 6 | DEE | NR | Infant (NR) | Epileptic spasms | global DD | chr20:47991279 A/AA p.Y274fsX36 | NA<br>NA<br>34 | not present | This report |
| 7 | DEE | Male | Toddler (1 yo) | Epileptic spasms, myoclonic, focal, GTC | global DD; severe ID | chr20:47991181 G/A p.R306C | Deleterious<br>Probably damaging<br>32 | not present | 8 |
| 8 | DEE | Female | Childhood (12 yo) | Atypical absence, myoclonic, GTC | global DD; moderate ID; ASD | chr20:47991181 G/A p.R306C | Deleterious<br>Probably damaging<br>32 | not present | This report |
| 9 | DEE | Female | Infant (6 mo) | Absence | Mild DD | chr20:47991181 G/A p.R306C | Deleterious<br>Probably damaging<br>32 | not present | This report |
| 10 | DEE | Female | Infant (6 mo) | Myoclonic, absence with eyelid myoclonia, GTC | global DD; mild-moderate ID | chr20:47991181 G/A p.R306C | Deleterious<br>Probably damaging<br>32 | not present | 16 |
| 11 | DEE | Female | Toddler (12-24 mo) | Absence, atonic, focal, myoclonic, tonic, GTC | global DD; ID | chr20:47991181 G/A p.R306C | Deleterious<br>Probably damaging<br>32 | Not present | This report |

| Case No. | Category | Sex | Developmental Stage at Seizure Onset (Age) | Seizure Types | Developmental Phenotype | Variant (hg19: NM_004975) | Prediction<br>SIFT<br>PolyPhen<br>CADD Score | gnomAD MAF | Ref |
| --- | --- | --- | --- | --- | --- | --- | --- | --- | --- |
| 12 | DEE | Male | Toddler (1 yo) | Absence, atonic, febrile, GTC | global DD; severe ID | chr20:47991181 G/A p.R306C | Deleterious<br>Probably damaging<br>32 | Not present | 11 |
| 13 | NDD | Male | NA | NR<br>(3 yo at last follow-up) | global DD | chr20:47991163 G/A p.R312C | Deleterious<br>Probably damaging<br>28.3 | not present | This report |
| 14 | NDD | Male | NA | NR<br>(6 yo at last follow-up) | global DD | chr20:47991163 G/A p.R312C | Deleterious<br>Probably damaging<br>28.3 | not present | This report |
| 15 | DEE | Male | Toddler (14 mo) | Focal, GTC | global DD; severe ID; ASD | chr20:47991162 C/T p.R312H | Deleterious<br>Probably damaging<br>32 | not present | 11 |
| 16 | DEE | Male | Infant (10 mo) | Epileptic spasms, myoclonic, atypical absence, | global DD; severe ID; ASD | chr20:47991162 C/T p.R312H | Deleterious<br>Probably damaging<br>32 | not present | 11 |
| 17 | DEE | Male | Toddler (18 mo) | NR; history of CSWS | global DD / ID | chr20:47991124 G/A p.R325W | Deleterious<br>Probably damaging<br>26.3 | not present | This report |
| 18 | NDD | Female | Childhood (4 yo) | One tonic seizure (5 yo at last follow-up) | global DD | chr20:47991107 C/G p.E330D | Deleterious<br>Probably damaging<br>22.8 | not present | This report |
| 19 | DEE | Female | Toddler (18 mo) | Focal, tonic, GTC | Global DD | chr20:47991107 C/G p.E330D | Deleterious<br>Probably damaging<br>22.8 | Not present | 13 |
| 20 | DEE | Male | Infant (9 mo) | Complex febrile, focal | global DD; moderate ID; ASD | chr20:47990989 A/G p.W370R | Deleterious<br>Probably damaging<br>29.6 | not present | This report |
| 21 | DEE | Male | Toddler (15 mo) | Myoclonic, tonic, atonic, GTC | Global DD; ID; ASD | chr20:47990964 A/G p.V378A | Deleterious<br>Probably damaging<br>25.9 | not present | This report |
| 22 | DEE | Female (deceased) | Toddler (13 mo) | Epileptic spasms, focal, GTC atonic, myoclonic, | global DD; ID | chr20:47990964 A/G p.V378A | Deleterious<br>Probably damaging<br>25.9 | not present | 9,11 |
| 23 | NDD | Female | NA | NR<br>(6 yo at last follow-up) | global DD; ID | chr20:47990965 C/G p.V378L | Deleterious<br>Probably damaging<br>26 | not present | 10 |
| 24 | DEE | Male | Infant (3 mo) | Epileptic spasms, tonic, GTC, myoclonic, atonic, absence | Global DD; profound ID | chr20:47990956 C/T p.G381R | Deleterious<br>Probably damaging<br>29.7 | not present | 11,12 |
| 25 | DEE | Male | Infant (8 mo) | Absence, myoclonic, tonic, GTC | Global DD | chr20:47990944 G/T<br>p.P385T | Deleterious<br>Probably damaging<br>27.6 | not present | This report |
| 26 | DEE | Male (deceased) | Toddler (13 mo) | Epileptic spasms, focal atonic, myoclonic, GTC | global DD; ID; ASD | chr20:47990944 G/T<br>p.P385T | Deleterious<br>Probably damaging<br>27.6 | not present | 11 |
| 27 | DEE | Female | NR | Seizures (not otherwise specified) | global DD | chr20:47990924 T/G<br>p.K391N | Deleterious<br>Probably damaging<br>23 | not present | 11,15 |
| 28 | DEE | Female | Toddler (14 mo) | GTC, reflex, atonic | global DD; profound ID; ASD | chr20:47990851 A/G<br>p.F416L | Deleterious<br>Probably damaging<br>28.9 | not present | 11,12 |
| 29 | DEE | Female | Childhood (3 yo) | Tonic, myoclonic, atypical absence | global DD; severe ID | chr20:47990851 A/G<br>p.F416L | Deleterious<br>Probably damaging<br>28.9 | not present | 11 |
| 30 | DEE | Female | Toddler (15 mo) | Epileptic spasms, myoclonic, absence, tonic, atonic | Global DD; severe ID | chr20:47990851 A/G<br>p.F416L | Deleterious<br>Probably damaging<br>28.9 | Not present | This report |
| 31 | DEE | Male | *(Inherited from unaffected parent)*<br>*Referred for whole exome sequencing* | | | chr20:47990726 G/C<br>p.S457R | Deleterious<br>Probably damaging<br>24.6 | $3.98 \times 10^{-6}$ | This report |
| 32 | DEE | Female | Infant (5 mo) | Epileptic spasms | gross motor delay | chr20:47990498 G/T<br>p.Y533x | NA<br>NA<br>36 | not present | 11 |

ASD: autism spectrum disorder; CSWS: continuous spike-wave during sleep; DD: developmental delay; DEE: developmental and epileptic encephalopathy; GTC: generalized tonic clonic; ID: intellectual disability; mo: months; NA: not applicable; NDD: neurodevelopmental disorder; NR: not reported; yo: year old.

**Figure 1.**
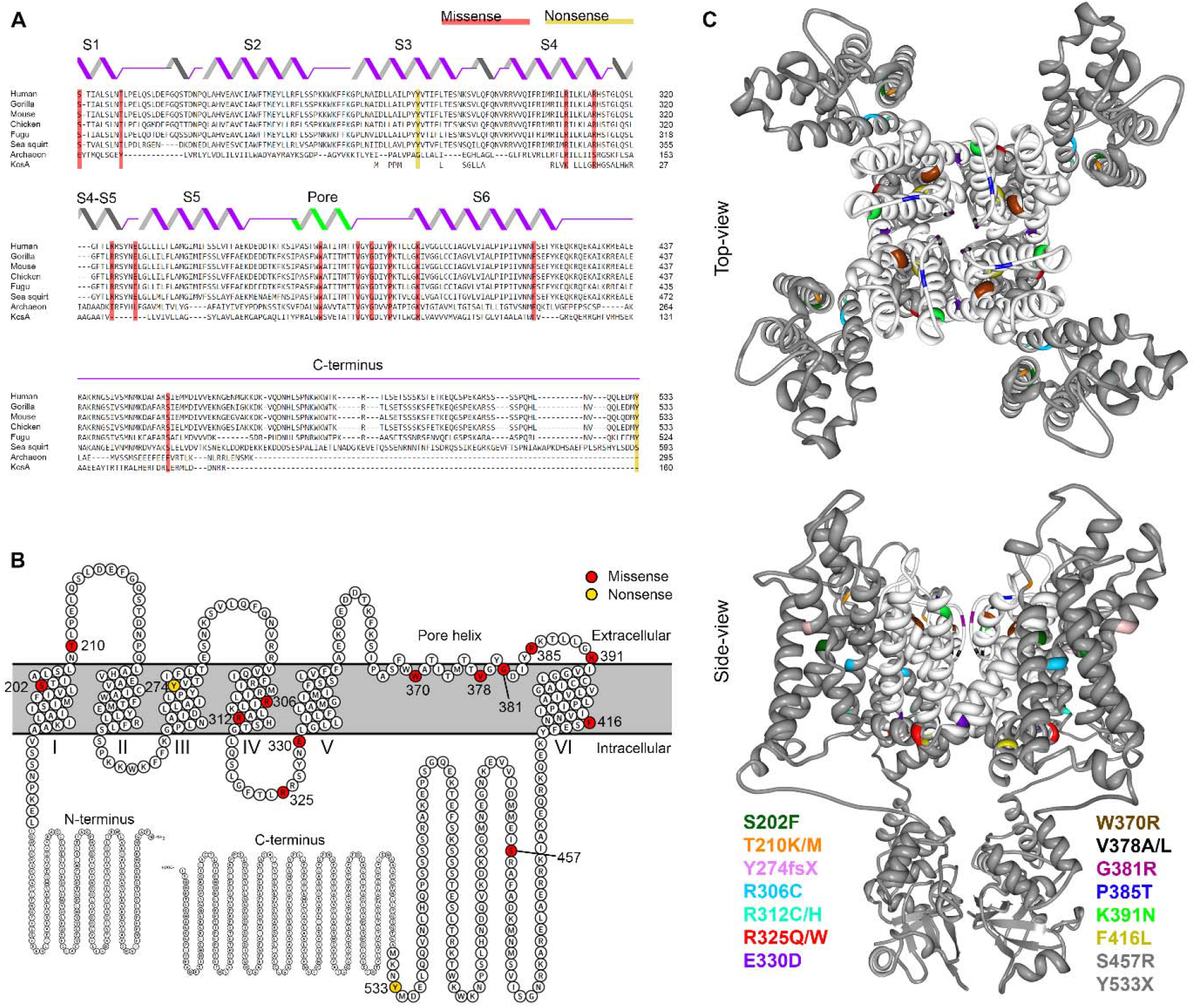
KCNB1 variants identified in individuals with epileptic encephalopathy. **A**. Evolutionary conservation of Kv2.1 shown by multiple sequence alignment of Kv2.1 species orthologues (Clustral Omega^41^); secondary structural elements are illustrated above the sequences. Kv2.1 variants are shaded in red (missense) and yellow (nonsense). **B**. Schematic view of the entire Kv2.1 subunit and the variants in the membrane: modified from Protter plot of Q14721 (KCNB1 _Human)^42^. **C**. Locations of variants (color-coded) mapped onto crystal structure of Kv2.1/Kv1.2 chimera (PDB2R9R)^23^: a top-view and a side-view across lipid bilayer. Pore domain is highlighted in white; two variants (S457R and Y533X) are not depicted, because their location in the distal C-terminus is not available in crystal structure.

Clinical phenotypes for the variants functionally characterized in this report are summarized in Table 1, and include DEE in 28 patients and NDD in four patients. Among those with NDD, one had a single seizure and three had no history of seizure at last reported follow-up at 6 years of age or younger. The phenotypes associated with novel variants reported in Table 1 are consistent with previous reports^7–14^ Table 1 also includes variants from previous clinically focused reports^11,16^, summarized here for completeness, with updates when available. In total, there are 17 pathogenic or likely pathogenic variants identified in 27 individuals. This includes a number of recurrent variants (T210M x3; R306C x6; R312C x2; R312H x2; E330D x 2; V378A x2; P385T x2; F416L x3), half of which occur at methyl-CpG dinucleotides that are known to be susceptible to deamination.

### Functional consequences of Kv2.1 variants: I_peak_ density

To screen for functional consequences of the missense or nonsense *KCNB1* variants, we performed automated patch clamp recording of these variants expressed in CHO-K1 cells. Wild-type (WT) or variant Kv2.1 channels were transiently transfected by electroporation and whole-cell currents recorded in voltage-clamp mode using the SyncroPatch 768PE automated patch clamp platform (Fig. 2A). Previous studies have shown the feasibility of this approach to study channel properties of Kv2.1 and the closely related Kv7.1 channel encoded by *KCNQ1*^14,27,28^. In addition, we also tested on this platform the T374I variant that we previously characterized by manual patch clamp recording^7^, and reproduced the loss-of-function phenotype (Fig. 2B; p < 0.001, unpaired t-test @ +60mV vs WT). This further validates use of this platform to screen for Kv2.1 dysfunction.

**Figure 2.**
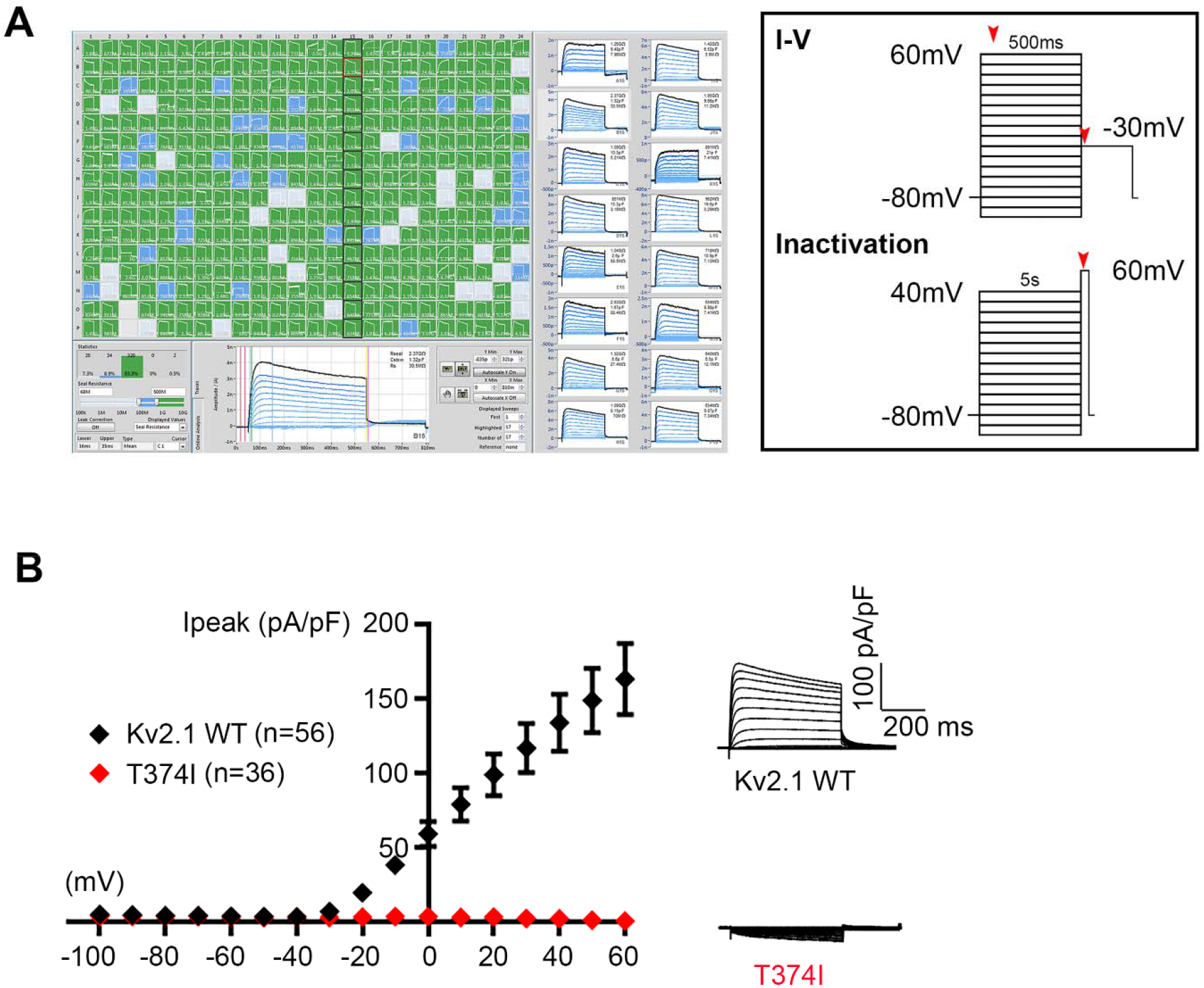
High-throughput functional screening of Kv2.1 variants. **A**. Screenshot of automated whole-cell current recordings from CHO-K1 cells transiently expressing Kv2.1 channels, with 16 individual cell recordings shown on the right blue traces. Voltage protocols used for functional studies are depicted in the box; red arrows indicate timepoints of current measurement. **B**. Validation of automated patch clamp recordings of Kv2.1 channels. Averaged Kv2.1 WT and T374I variant, (previously published by our group using manual patch clamp) current-voltage relationships and whole-cell current density traces show a similar loss-of-function phenotype for the T374I variant.

The Kv2.1 channel is composed of four alpha subunits encoded by *KCNB1* that form a tetrameric assembly. When singly expressed in CHO cells, the Kv2.1 channels assume a homomeric configuration consisting of 4 identical WT or variant subunits. In the homomeric configuration, the majority of the Kv2.1 variants (13/19) exhibited low K^+^ conductance (loss-of-function) (Fig. 3A, red bars), while the remaining variants displayed moderate K^+^ conductance (partial loss-of-function) (blue bars), or K^+^ conductance similar to WT (grey bars). The degree of loss-of-function was considered ‘partial’ if more than 50% reduction and ‘severe’ if more than 75% reduction in peak current density compared to WT. The averaged whole-cell current density traces for Kv2.1 WT and representative variants are displayed to show the diversity in peak current density (Fig. 3A right). Averaged whole-cell current density traces for all Kv2.1 variants are shown in supplementary Figure 1. Averaged endogenous currents recorded from non-transfected CHO-K1 cells were digitally subtracted.

**Figure 3.**
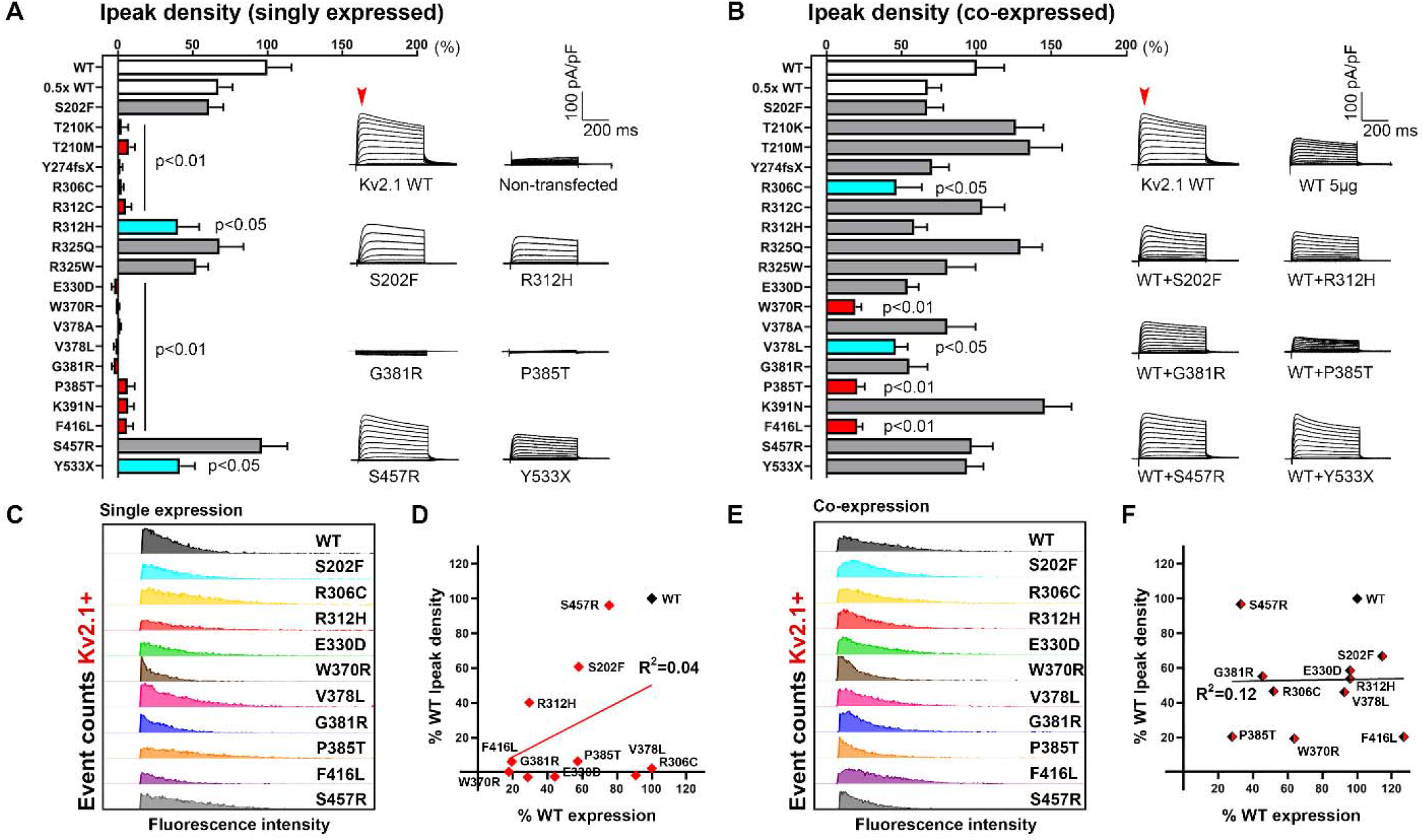
High-throughput functional screening of Kv2.1 variants. **A-B**. Comparison of peak current density of Kv2.1 variants in homomeric and co-expression (with WT) configuration (red bars p<0.001 vs WT, p<0.05 vs 0.5x WT; turquoise bars p<0.05 vs WT; One-way ANOVA Dunnett’s multiple comparisons). Averaged whole-cell current traces of selected variants are shown on the right; endogenous current recorded from non-transfected CHO-K1 cells were subtracted off-line. Red arrow denotes the time of peak current measurement. For co-expression configuration, Kv2.1 variants were co-expressed with WT by electroporation of equimolar amount of DNA. **C&E**. Protein expression analyses of variants by flow cytometry in the homomeric (left) and co-expression (right) configuration confirm Kv2.1 protein expression in variant-expressing cells, in particular for the ones with severely reduced current density, albeit with differing degrees of severity. Data are shown in histograms aligned to correct for between-run variability in fluorescence intensity; Y-axes were uniformly set at a maximum of 250 and 350 event counts (for C and E respectively) across conditions. **D&F**. Correlation of peak current density and protein expression by percent-Kv2.1 positive cells, in either homomeric or co-expression configuration, did not reach statistical significance (p=0.22 and 0.96 in order; Pearson’s correlation).

*KCNB1* DEE is associated with *de novo* heterozygous variants. Thus, an individual has Kv2.1 subunits produced from both WT and variant alleles. In order to determine the effect of the *KCNB1* variants in a condition that resembles heterozygosity in the patient, we co-expressed each variant along with wild-type channels in a 1:1 equimolar ratio in CHO-K1 cells. In the co-expression configuration (WT+variant), many variants exhibited peak current density similar to cells transfected with WT+WT channels, suggesting that these variants may be rescued by WT co-expression. In contrast, the W370R, P385T and F416L variants exhibited significantly lower peak current density when compared to WT+WT cells (Fig. 3B, bars in red). These variants also showed significantly lower peak current density when compared to cells transfected with half the amount of WT plasmid (0.5x WT), and suggest a dominant-negative interaction of the variant with WT subunits. The R306C and V378L variants exhibited a partial loss-of-function (bars in turquoise). Averaged current traces for a subset of variants are displayed (Fig. 3B right). Complete statistical information including number of cells and p-values is listed in Supplementary Table 1.

### Total protein expression of Kv2.1 variants

To test whether loss-of-function phenotypes, defined by impaired K^+^ current density, was due to absence or reduction of channel protein, we evaluated total protein expression levels for Kv2.1 variants using immunoblotting and/or immunocytochemistry-flow cytometry. Frozen aliquots of the same cells tested by electrophysiology above were used for this experiment. In the homomeric configuration, population analyses of Kv2.1-positive cells confirmed that *KCNB1* variants with severe loss-of-function phenotypes had lower levels of Kv2.1 protein compared to WT (Fig. 3C&E). The same variants when co-expressed 1:1 with WT showed similar results. Correlational analyses of peak current density and total protein expression of the variants showed no relationship between parameters in either the homomeric or coexpression condition (Fig. 3D&F; in red, R^2^= 0.16; p=0.22, and in black, R^2^= 0.0003; p=0.96 respectively; Pearson’s). Variants not shown in Fig. 3C&E were tested with western blot analyses (Supplementary Fig. 2A-C).

### Voltage-dependence of channel activation and inactivation

Voltage-dependence of channel activation and inactivation was assessed for the subset of Kv2.1 variants with sufficient K^+^ conductance to reliably measure these parameters. In the homomeric configuration, a subset of variants with K^+^ conductance exhibited altered voltage-dependence of channel activation and inactivation when compared to the WT channel (Table 2). The S202F and R312H variants exhibited substantial depolarizing shifts in their V_1/2_ for both channel activation and inactivation (Fig. 4A) and significantly larger slope factors. This suggests a potential decrease in the sensitivity of channel inactivation to voltage change. The Y533X variant also induced a depolarizing shift in V_1/2_ of activation without affecting channel inactivation (Table 2). In contrast, the R325W variant caused hyperpolarizing shifts in the V_1/2_ values for both channel activation and inactivation (Fig. 4A).

**Table 2.** Biophysical properties of Kv2.1 variants singly or co-expressed with WT. Singly expressed variants with at least 10pA/pF are included. Values that differ from the respective WT condition are shown in red, with asterisks denoting significance levels as *p<0.05, **p<0.01, ***p<0.001.

| Configurations<br>or<br>variants | Voltage-dependence of activation |  | Voltage-dependence of inactivation |  |
| --- | --- | --- | --- | --- |
|  | V<sub>1/2</sub> | k | V<sub>1/2</sub> | k |
| **WT** | -4.6 ± 1.0 | 13.0 ± 0.4 | -35.5 ± 2.6 | 7.5 ± 0.6 |
| **S202F** | 24.3 ± 3.8<br>n=14*** | 16.4 ± 1.7<br>n=14 | 2.7 ± 1.2<br>n=30*** | 8.8 ± 0.4<br>n=30*** |
| **R312H** | 28.1 ± 1.5<br>n=21*** | 18.2 ± 1.0<br>n=21** | -0.1 ± 2.3<br>n=18*** | 9.9 ± 0.7<br>n=11*** |
| **R325Q** | -5.0 ± 2.5<br>n=18 | 14.8 ± 1.2<br>n=18 | -39.6 ± 1.8<br>n=17 | 7.1 ± 0.4<br>n=23 |
| **R325W** | -23.4 ± 3.9<br>n=22*** | 17.7 ± 1.2<br>n=22* | -58.0 ± 1.4<br>n=25*** | 5.7 ± 0.2<br>n=25*** |
| **S457R** | -13.3 ± 1.4<br>n=25 | 12.0 ± 0.7<br>n=25 | -44.2 ± 1.7<br>n=38 | 6.5 ± 0.2<br>n=38 |
| **Y533X** | 13.0 ± 3.6<br>n=17*** | 18.3 ± 1.4<br>n=17*** | -45.7 ± 2.6<br>n=26 | 6.8 ± 0.6<br>n=26 |
| **WT+WT** | -7.3 ± 0.9 | 12.9 ± 0.9 | -39.6 ± 0.6 | 6.3 ± 0.1 |
| **WT+S202F** | 1.3 ± 3.1<br>n=25 | 22.2 ± 0.8<br>n=25 *** | -29.6 ± 1.7<br>n=54** | 10.4 ± 0.4<br>n=54*** |
| **WT+T210K** | -10.8 ± 1.3<br>n=44 | 13.1 ± 0.5<br>n=44 | -46.2 ± 1.3<br>n=34** | 6.3 ± 0.2<br>n=34 |
| **WT+T210M** | -6.9 ± 1.7<br>n=29 | 12.9 ± 0.9<br>n=29 | -43.3 ± 2.4<br>n=25 | 6.5 ± 0.3<br>n=25 |
| **WT+Y274fsX** | -7.1 ± 4.0<br>n=18 | 14.5 ± 1.1<br>n=18 | -40.2 ± 1.6<br>n=65 | 5.7 ± 0.2<br>n=65 |
| **WT+R306C** | -15.9 ± 3.7<br>n=11 | 15.3 ± 2.2<br>n=11 | -44.9 ± 3.2<br>n=22 | 9.0 ± 0.6<br>n=22** |
| **WT+R312C** | -9.1 ± 1.3<br>n=35 | 13.4 ± 0.5<br>n=35 | -42.0 ± 1.3<br>n=27 | 6.0 ± 0.1<br>n=27 |
| **WT+R312H** | 0.1 ± 2.7<br>n=31 | 14.3 ± 0.7<br>n=31* | -38.7 ± 1.5<br>n=78 | 7.7 ± 0.4<br>n=78* |
| **WT+R325Q** | -10.7 ± 1.4<br>n=43 | 12.8 ± 0.6<br>n=43 | -43.4 ± 1.7<br>n=32 | 5.9 ± 0.2<br>n=32 |
| **WT+R325W** | -14.3 ± 1.7<br>n=39* | 16.1 ± 1.0<br>n=39 | -51.9 ± 1.5<br>n=35*** | 5.9 ± 0.2<br>n=35 |
| **WT+E330D** | -12.4 ± 2.3<br>n=18 | 15.3 ± 1.4<br>n=18 | -40.4 ± 2.3<br>n=52 | 7.4 ± 1.0<br>n=52 |
| **WT+W370R** | -3.5 ± 3.3<br>n=12 | 16.4 ± 1.8<br>n=12 | -38.9 ± 2.9<br>n=26 | 6.9 ± 0.4<br>n=26 |
| **WT+V378A** | -7.5 ± 1.4<br>n=29 | 13.3 ± 0.5<br>n=29 | -41.4 ± 1.2<br>n=32 | 5.6 ± 0.2<br>n=32 |
| **WT+V378L** | -5.8 ± 4.8<br>n=20 | 16.0 ± 1.2<br>n=20 | -46.4 ± 1.9<br>n=61* | 7.5 ± 0.3<br>n=61** |
| **WT+G381R** | -9.4 ± 2.7<br>n=27 | 14.2 ± 1.0<br>n=27 | -41.2 ± 1.4<br>n=62 | 6.3 ± 0.5<br>n=62 |
| **WT+P385T** | -6.7 ± 4.1<br>n=11 | 14.9 ± 1.2<br>n=11 | -40.6 ± 2.1<br>n=43 | 6.9 ± 0.5<br>n=43 |
| **WT+K391N** | -9.4 ± 1.2<br>n=44 | 12.6 ± 0.4<br>n=44 | -41.4 ± 1.2<br>n=39 | 5.9 ± 0.1<br>n=39 |
| **WT+F416L** | -3.1 ± 3.7<br>n=16 | 15.6 ± 1.1<br>n=16 | -39.5 ± 1.6<br>n=48 | 7.3 ± 0.4<br>n=48 |
| **WT+S457R** | -18.9 ± 3.0<br>n=22* | 14.1 ± 1.0<br>n=22 | -42.0 ± 1.6<br>n=42 | 5.9 ± 0.2<br>n=42 |
| **WT+Y533X** | -11.6 ± 1.7<br>n=42 | 14.8 ± 0.8<br>n=57 | -43.0 ± 1.4<br>n=65 | 6.0 ± 0.2<br>n=65 |

**Figure 4.**
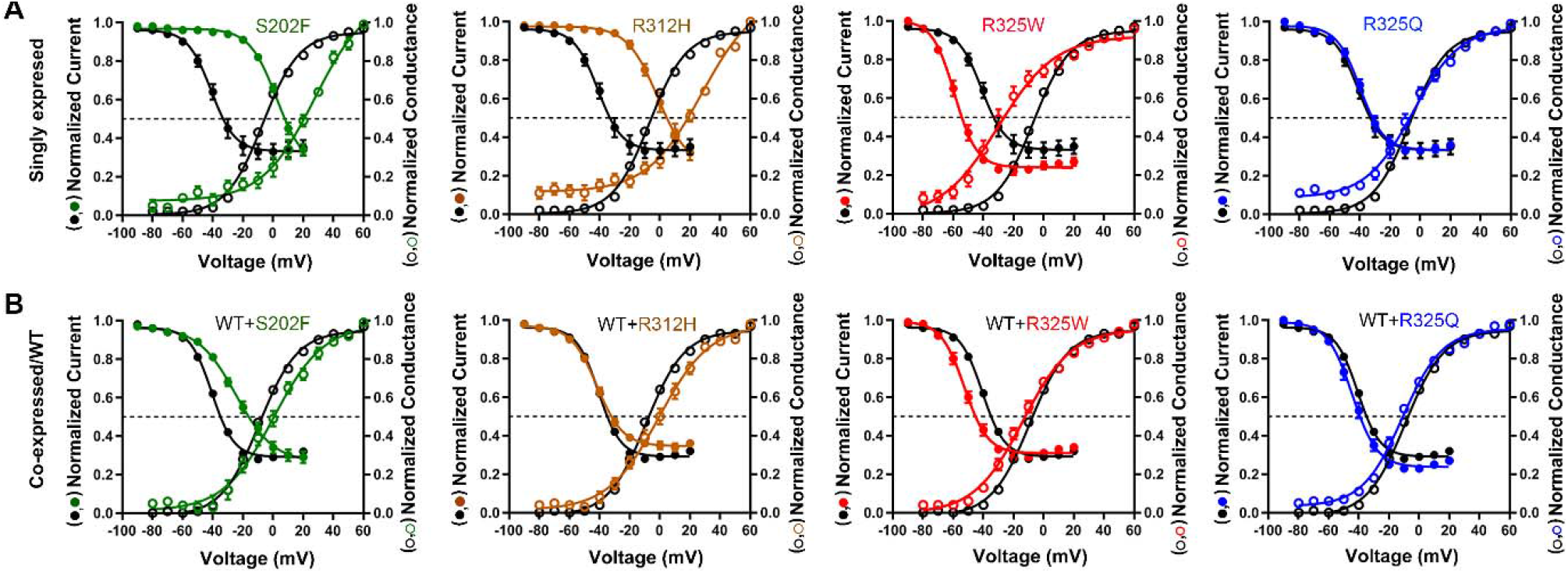
Voltage-dependence of activation/inactivation of Kv2.1 variants. **A-B.** Voltage-dependent channel activation (open circles) and inactivation (solid circles) curves obtained from selected homomeric and co-expression Kv2.1 variants, compared to WT (circles in black). Solid lines represent averaged data fits with the Boltzmann equation. Numerical values of V_1/2_ and slope factor (*k*) for the other variants are listed in Table 2.

When co-expressed with WT, the effects of R312H on channel activation and inactivation were abolished, suggesting rescue by WT subunits. Co-expression with WT channels prevented the depolarizing shift in activation due to S202F. However, the observed differences in V_1/2_ and *k* inactivation for homomeric S202F channels were still present when co-expressed with WT channels (Table 2). The hyperpolarizing shifts in V_1/2_ for activation and inactivation caused by the R325W mutation were still present in the presence of WT subunits, albeit with smaller magnitude. Although the T210K variant was non-functional when expressed as homotetramer, it caused a hyperpolarizing shift in inactivation V_1/2_ when co-expressed with WT. Lastly, co-expression of WT and S457R channels resulted in a hyperpolarizing shift in V_1/2_ that was not observed for homomeric S457R channels (Table 2). A complete statistical summary including V_1/2_ value and slope factor *k* for all variants is presented in Table 2.

### Functional effects of non-pathogenic variants

Two likely non-pathogenic variants (R325Q and S457R) were included in the functional assays, with the experimenters blinded to this status. The R325Q variant was reported as a singleton variant (MAF 3.983^e-06^) in the genome aggregation database (gnomAD) that is devoid of individuals with severe pediatric disease, as well as their first-degree relatives^21^. Unlike the disease-associated R325W mutant, which induced changes in both channel activation and inactivation, the R325Q variant did not exhibit any significant changes in the properties analyzed, and thus behaved like the WT channel (Fig. 4A-B).

Another variant S457R was classified as non-pathogenic based on inheritance from an unaffected parent and presence of this S457R as a singleton variant (MAF 3.983^e-06^) in gnomAD^21^. In the homomeric configuration, S457R did not differ from WT on any measured parameters, although there was a hyperpolarizing shift in the voltage-dependence of activation in the co-expressed configuration. S457 is subject to post-translational modification by phosphorylation, and dephosphorylation of Kv2.1 is associated with hyperpolarizing shifts in voltage-dependent gating.^29^ Thus, the observed difference in voltage-dependence of activation may reflect differences in phosphorylation state between the mutant (i.e. non-phosphorylated residue 457) and WT subunits (i.e. phosphorylated residue 457) under our experimental conditions, which require intracellular sodium fluoride, a known phosphatase inhibitor, for gigaohm seal formation.

### Expression analysis of Kv2.1 variants: cell-surface trafficking

Presence at the cell-surface is critical for voltage-gated ion channel function, and some variants may interfere with efficient cell-surface expression of Kv2.1 protein. Cell-surface and total expression levels of Kv2.1 protein were simultaneously examined using immunocytochemistry followed by flow cytometry (FC). An extracellular HA epitope tag enabled discrimination of cell surface and total Kv2.1 fractions allowing for rapid detection of pathogenic variants with deficient cell-surface expression. A substantial population of WT Kv2.1 reached the cell-surface, with a mean surface/total expression ratio higher than 0.8, indicating highly efficient trafficking of Kv2.1 to the cell-surface (Fig. 3A-B), as previously shown for Kv2.1 in other experimental models^30,31^. The majority of the variants (13/19) exhibited pronounced deficits in cell-surface expression of Kv2.1 protein (Fig. 5A-B), potentially explaining the loss of K^+^ conductance observed from high-throughput electrophysiology screening (Fig. 3C). The non-pathogenic variants, R325Q and S457R, exhibited cell-surface and total expression that was indistinguishable from WT (Fig. 5A-C). Cross-validation of select variants (T210K/M, R312C, R325Q/W, V378A, and K391N) with cell-surface biotinylation followed by immunoblotting produced congruent results (Supplementary Fig. 2A-C).

**Figure 5.**
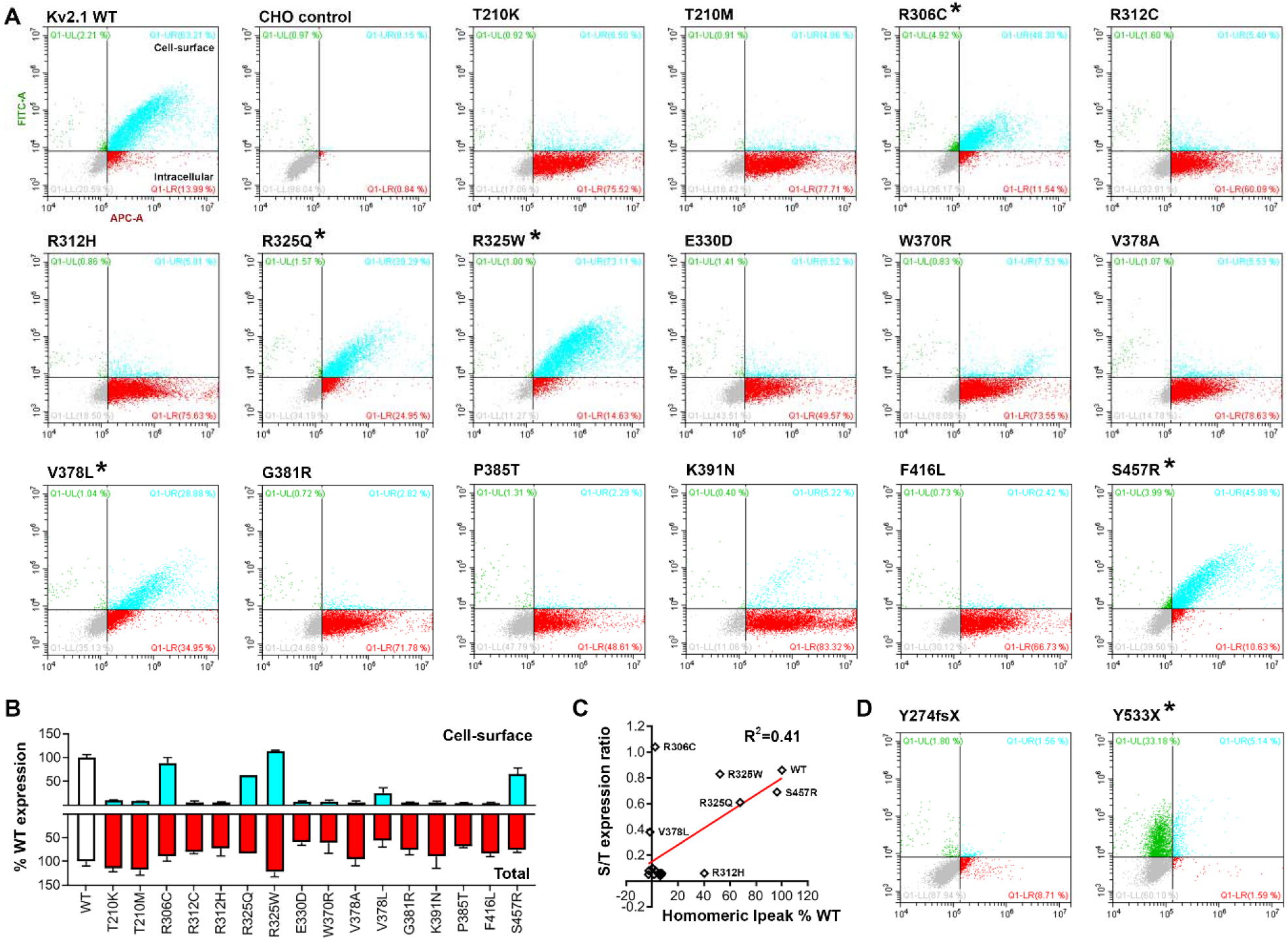
Total and cell surface protein expression levels of Kv2.1 variants. **A**. Representative dot-plots of flow cytometry analyses of CHO cells expressing Kv2.1 tagged with HA-epitope at the extracellular site between S1-S2 linker regions; anti-HA-Alexa488 and anti-Kv2.1-Alexa647 antibodies were used to probe the cell-surface and intracellular Kv2.1 respectively. Population in turquoise (HA^+^/Kv2.1^+^) denote Kv2.1 located at the cell-surface, while the dots in red (HA+/Kv2.1^−^) indicate expressed Kv2.1 protein located intracellular. WT and non-transfected CHO cells (negative control) are shown at the top left for comparison. Nontransfected CHO cells underwent the entire immunocytochemistry labeling protocol. **B**. Expression level comparisons of Kv2.1 variants to that of WT (white bars) revealed reduced cell-surface trafficking of variants (* denote variants with intact cell-surface trafficking, i.e. S/T ratio > 0.3): cell-surface expression in turquoise and the total expression in red. **C**. Correlational analyses of surface-to-total expression level ratio and homomeric Ipeak density (values in % WT) result in R^2^=0.41 and p=0.006: Pearson’s. **D**. Analyses of two truncation variants, devoid of the epitope site for anti-Kv2.1-Alexa647, further validate the assay. The Y274fsX36 variant fails to reach the cell-surface. In contrast, for the Y533X, the population in green (HA+/Kv2.1^−^) denote Kv2.1 expressed and reaching the cell-surface, yet unable to be detected by the anti-Kv2.1-Alexa647 antibody due to the absent epitope.

The S202F variant located near the HA-epitope insertion site (a.a.220) likely had altered immunoreactivity considering the lack of anti-HA signal but intact peak current density of S202F. It is possible that S202F results in steric hindrance caused by the substituted phenylalanine, preventing the HA antigen-antibody interaction (Supplementary Fig. 2D). For the T210K/M variants, also located nearby the HA-tag insertion site, immunoblot analyses of the non-tagged T210K/M variants confirmed the reduced cell-surface expression by flow cytometry (Supplementary Fig. 2 and Fig. 5A-B), which is consistent with their loss-of-function phenotypes in homomeric configuration (Fig. 3A).

The two truncation variants, Y274fsX36 and Y533X, were also tested for expression level analyses, despite lacking the epitope site for anti-Kv2.1-Alexa647 (a.a. 837-853 at C-terminus). The Y274fsX variant did not have cell-surface HA antigenicity. However, in a subsequent experiment, permeabilization prior to anti-HA incubation revealed abundant total expression, suggesting a trafficking defect for the truncated Y274fsX36 protein. The Y533X variant resulted in the cell population that showed HA^+^/Kv2.1^−^ immunoreactivity (Fig. 5D right), indicating that truncated Y533X proteins reach the cell-surface.

### Multivariate transformational analyses of Kv2.1 variants

To visualize the relationship among various parameters measured in this study, we performed principal component analyses using a total of 6 variables measured for the homomeric configuration: Ipeak density, V_1/2_ and *k* for activation and inactivation, protein expression cell-surface/total ratio (Fig. 6). The top 3 leading dimensions for PC1 were Ipeak density (0.61), cell-surface protein/total ratio (0.62), and V_1/2_ for activation (−0.28); and for PC2 were Ipeak density (0.62), and inactivation V_1/2_ (0.47) and k (0.49). Plotting PC1 vs. PC2 separated the nonpathogenic group (R325Q/S457R and WT; in green) from the pathogenic variants with 95% confidence. However, it was not possible to separate the pathogenic variants by clinical classification.

**Figure 6.**
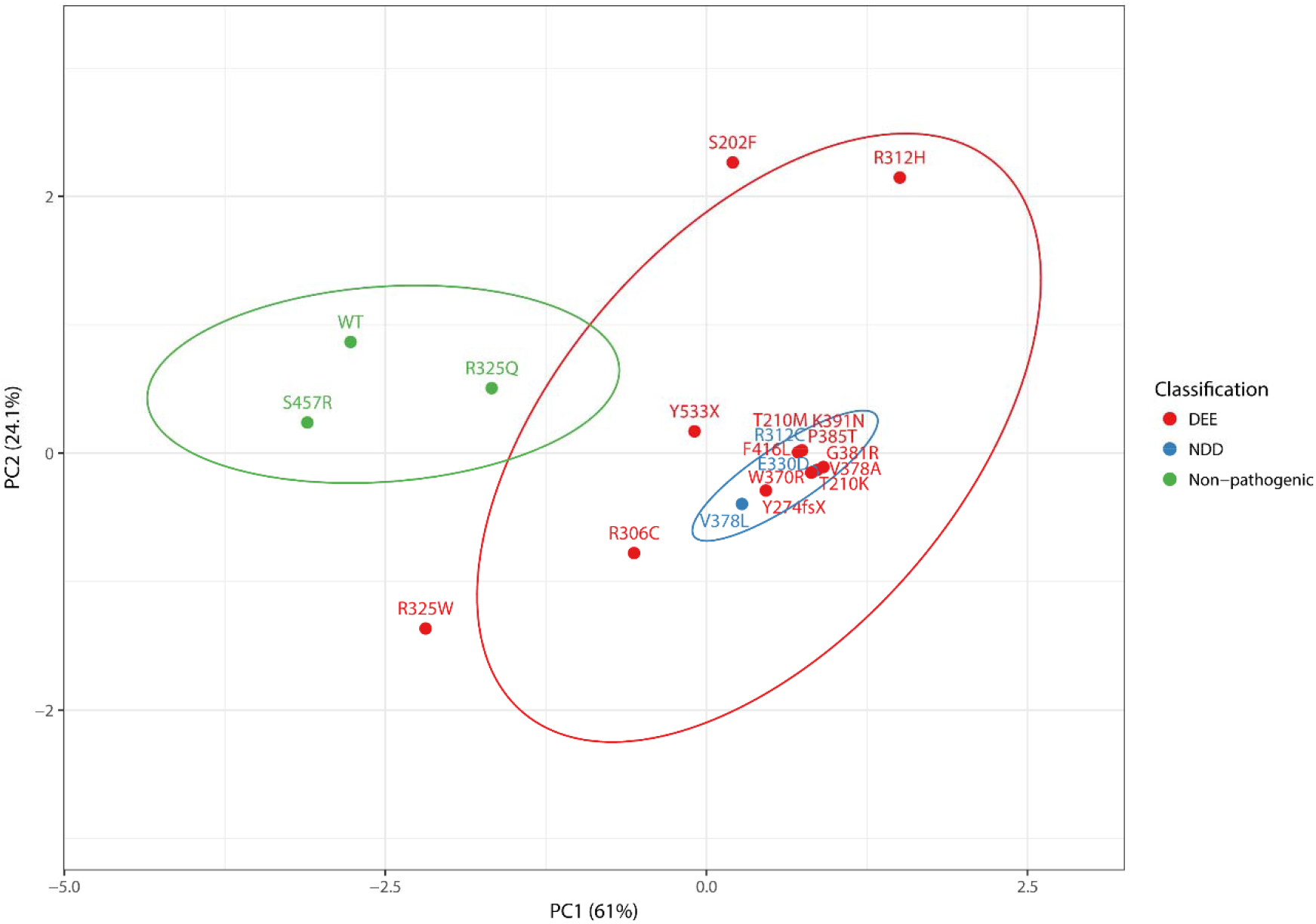
Principal component analyses of Kv2.1 variants. Experimental data for biophysical and protein expression analyses were transformed with respect to WT, to generate eigenvector-based multivariate analyses, and plotted using ClustVis^20^. Nipals PCA is used to calculate principal components, with ellipses depicting 95% confidence interval. Each axes explains 61% and 24.1% of the total variance (X and Y, in order): N = 20 data points. Total of 6 variables measured in the homomeric condition were used: Ipeak density, V_1/2_ and *k* for activation and inactivation, protein expression cell-surface/total ratio. The top 3 leading dimensions (variable) for PC1 were Ipeak density (0.61), cell-surface/total expression ratio (0.62), V_1/2_ for activation (−0.28); and for PC2 were Ipeak density (0.62), and inactivation V_1/2_ (0.47) and *k* (0.49). Nonpathogenic variants and WT (in green) are separable from the pathogenic variants (red, blue). DEE, developmental epileptic encephalopathy; NDD, neurodevelopmental disorder.

## DISCUSSION

### Kv2.1 loss-of-function in *KCNB1* DEE

Here we report high-throughput functional studies of 19 *KCNB1* variants, including 17 likely pathogenic variants identified in individuals who underwent clinical genetic testing for epileptic encephalopathy and/or global DD or ID of unknown cause, as well as two presumed non-pathogenic variants present in the gnomAD population database that excludes individuals with severe pediatric disease^21^. High-throughput electrophysiological and biochemical analysis of Kv2.1 currents in cells transiently expressing these variants revealed partial or near-complete loss-of-function as a common feature, consistent with previous reports using manual electrophysiology on a limited number of *KCNB1* variants^7–9^. The molecular mechanisms underlying loss-of-function phenotypes included lower peak current density due to low total and/or cell surface protein expression, as well as shifts in voltage-dependence of activation and inactivation.

The majority of variants exhibited significantly lower peak current density when expressed as homotetramers. However, for many of those variants, co-expression with WT subunits normalized peak current density. Rescue by the presence of WT subunits suggest that variant subunits traffic to the cell surface as part of a heteromeric assembly, which raises the possibility for phenotypic rescue by molecular chaperones. In contrast, co-expression of some variants (R306C, V378L) with WT resulted in lower current density that was similar in magnitude to 0.5x WT, suggesting that while the mutant subunits cannot be rescued functionally, they do not interfere with cell surface expression of WT. Other variants (W370R, P385T, F416L) when co-expressed with WT exhibited lower current density than 0.5x WT, suggesting a potential dominant negative effect on cell surface expression of WT subunits.

Several variants encoded functional Kv2.1 channels with current density indistinguishable from WT in the homotetramer configuration, but exhibited defects in voltage-dependence. Depolarizing shifts in V_1/2_ for both activation and inactivation were observed for S202F and R312H, resulting in loss-of-function due to impaired channel opening in a physiological voltage range. Conversely, R325W had hyperpolarized shifts in V_1/2_ for activation and inactivation. Although this likely results in enhanced channel opening at physiological voltages, the corresponding shift in the voltage-dependence of inactivation limits channel availability in the voltage range where Kv2.1 normally operates, suggesting overall loss-of-function. Co-expression of WT channels to approximate heterozygosity resulted in attenuated or altered voltage-dependence. This may be explained by variable populations of WT+WT, WT+var, and var+var channels. The presence of significant voltage-dependence phenotypes that were specific to co-expression conditions (i.e. activation *k* for WT+S202F or activation V_1/2_ for WT+S457R) suggests that for at least some variants, WT and variant subunits co-assemble in heterotetrameric channels.

### Comparison of same site variable substitutions

The importance of experimental validation for individual variants is highlighted by pairwise comparison of substitutions of different amino acids at the same position (i.e. T210K/M, R312C/H, R325Q/W and V378A/L). The T210K/M (CADD score of 28.1 and 28.2, respectively) variants resulted in severe loss of K^+^ conductance and cell-surface Kv2.1 expression. Threonine 210 has been shown to be a critical residue forming the S1-pore interface and mutation of the corresponding position in other Kv channels resulted in no detectable current^32,33^. A hydroxyl group has been shown to be required at this position for channel function^33^. Thus, substitution with either lysine or methionine would be incompatible with S1-pore coupling and channel function. In contrast, R312H and R312C resulted in partial and severe loss-of-conductance, respectively, which was the opposite of the *in silico* consequence prediction that suggested R312H (CADD score of 32) would be more deleterious than R312C (CADD score of 28.3). Retaining a positive charge at this critical position in the S4 segment by substituting histidine may lessen severity relative to a charge-neutralization cysteine substitution. Similarly, the R325Q/W variants differed in their voltage-dependence phenotypes, with the R325Q retaining WT-like behavior, consistent with non-pathogenicity. This suggests that substitution of arginine 325 with a bulky hydrophobic tryptophan^34^ may alter electrochemical coupling between S4-S5 and the S6 segment, while substitution with glutamine at this position is tolerated.

Both the V378A/L variants resulted in loss of K^+^ conductance in the absence of WT subunits. Protein expression analyses showed modest expression of V378L at the cell-surface, while the V378A variant was absent. This indicates discrete mechanisms underlying the loss of K^+^ conductance for these two variants. Leucine is present at the equivalent position in Kv3 and Kv4 channels, indicating that this substitution is structurally tolerable and maintains K^+^ selectivity, although it likely alters single channel conductance^35,36^. Conversely, substitution with the smaller alanine has been shown to disrupt ion selectivity and may interfere with Kv2.1 channel stability^36^. Previous studies of the V378A variant in CHO-K1 and COS-7 cells showed impaired expression that could be rescued by application of the Kv2 inhibitor GxTx, suggesting enhanced internalization and degradation as a possible mechanism underlying deficient expression^9^. Consistent with this, we observed robust total expression of V378A at 24 hours post-transfection by immunocytochemistry/flow-cytometry (Fig. 5), while immunoblotting at 72 hours showed little V378A expression (Fig S2).

### High-throughput variant characterization pipeline

One of the goals of this study was to leverage new electroporation and automated planar patch clamp technologies to develop a high-throughput pipeline for functional and biochemical characterization of *KCNB1* variants identified by clinical genetic testing. With this optimized pipeline, it is feasible to functionally characterize new variants rapidly. Moreover, the immunocytochemistry-flow-cytometry assay developed here enables high-throughput detection of cell-surface and total protein expression levels for further delineation of the mechanisms by which a variant affects channel function. Efficient performance of functional and biochemical studies of ion channel variants is a critical step to realize the precision medicine goal of rapid, definitive precision diagnosis and eventually targeted treatment based on the individual’s specific genetic variant.

### Utility of functional characterization for clinical genetic testing

Based on ACMG guidelines, evidence of a deleterious effect on protein function can be considered as ‘strong’ evidence for pathogenicity, given that the functional assay is robust and validated^3^. Electrophysiology is considered the gold standard for characterizing the effects of ion channel variants, and is a well-validated, robust approach. In contrast, *in silico* evidence based on pathogenicity prediction algorithms such as CADD, M-CAP, or others are only considered ‘supporting’ evidence, thus insufficient when considered alone. Functional variant characterization, both in channelopathies and other disorders, has untapped potential to provide significant diagnostic benefit for determining whether a particular variant is likely pathogenic. Our goal with *KCNB1* is to functionally characterize a large series of variants to serve as a reference dataset for clinical genetic test interpretation in newly diagnosed patients, and to provide additional evidence for potential re-classification of variants previously classified as variants of uncertain significance.

The *KCNB1* variants R325Q and S457R described herein provide a concrete example of how functional variant characterization can provide useful diagnostic information. R325Q is a rare population variant that exhibited similar functional characteristics to WT channels, suggesting that arginine 325 is a residue tolerant to substitution with glutamine (Q). However, R325W has significant functional defects both as a homotetramer and when co-expressed with WT (Fig. 3A&B), suggesting this variant is likely pathogenic. Further strengthening the case for functional characterization, *in silico* pathogenicity predictions such as CADD scores actually predicted the opposite, with R325Q predicted as pathogenic (CADD=20.2) and R325W as likely benign (CADD=15.8). Another example concerns S457R, which was initially identified by clinical genetic testing in a child with DD and epilepsy and was later found upon parental testing to be inherited from an unaffected parent. Prior to parental testing, we performed an electrophysiological study of S457R in the homomeric configuration and found no deleterious effect on protein function, suggesting that this variant was non-pathogenic. Principal component analysis using measured parameters from the functional studies allowed reliable separation of nonpathogenic group, consisting of R325Q and S457R variants along with the WT, from the other 17 variants studied, highlighting the viability of high-throughput functional screening to differentiate nonpathogenic from pathogenic variants with high confidence (95%).

### Limitations of high-throughput functional screening

Despite the aforementioned strengths of high-throughput screening of genetic variants in a heterologous expression model, there are some limitations. Some of the Kv2.1 variant effects detected in our study may become less or more pronounced in neurons, as neurons are highly specialized cells with diverse distinct morphologies compared to CHO cells. For example, variant and WT alleles are co-expressed in individuals with *KCNB1* DEE, which is associated with heterozygous variants. However, in the heterologous expression system, co-expression with WT sometimes masked variant effects. This highlights a difference between heterologous expression versus the nervous system, and suggests that homomeric expression provides a more robust assay. In addition, conducting K^+^ is not the only function of Kv2.1 in neurons, as a population of non-conducting Kv2.1 clustered at plasma membrane-endoplasmic reticulum junctions operate as a docking site for membrane proteins^31^. Lastly, for variants that alter binding site affinity for interacting proteins like KChAP, AMIGO, and KCNE1-3^37–39^, their effect may not be detected if these proteins are not present at physiological levels in this model. Hence, additional characterization of variants in more complex experimental systems such as knock-in mice or patient-derived iPSCs may be needed for more complete understanding of disease mechanisms in neurons or other cells where Kv2.1 is a significant contributor (e.g., pancreatic beta cells).

Another goal of our study was to assess potential correlations between Kv2.1 channel dysfunction and clinical severity. We used principal component analysis to address this question using 6 parameters that were measured experimentally. Although the clinical phenotypes of DEE and NDD were not distinguished, it is possible that the NDD cases (3-6 years of age at the time of reporting) will go on to develop epilepsy, as there are at least two DEE cases with seizure onset in adolescence (Table 1). Alternatively, the underlying pathogenic mechanisms may require more complex experimental systems like neurons or *in vivo* animal models, and/or that final expression of the clinical phenotype can be influenced by other genetic, epigenetic or environmental modifiers.

### Conclusions

In summary, many of the disease-associated *KCNB1* variants (14/17) we studied exhibit loss-of-function phenotypes that ultimately result in a channel hypofunction, through 1) decreased K^+^ conductance, 2) altered voltage-dependence of channel activation or inactivation, 3) reduced protein expression, or 4) altered cell-surface trafficking. Classification of *KCNB1* variants into different categories of defects indicates that molecular chaperones to increase cell-surface expression may be a suitable therapeutic strategy for some patients. A subtype-selective activator of Kv2.1 may be a viable therapy for others, although this would not address non-conducting functions of Kv2.1^31,40^, which may contribute to disease pathophysiology. Alternatively, gene therapy approaches that increase expression of the WT allele may hold promise for treatment of patients with *KCNB1* variants. It is thus critical to be able to efficiently and reliably characterize *KCNB1* variants in patients with DEE or DD/ID as pathogenic to provide the precision diagnoses required for precision treatment.

## ACKNOWLEDGEMENTS

We thank the patients and their families for their cooperation, and Tatiana Abramova, Reshma Desai, Nicole Hawkins, Tyler Thaxton, and Alex Huffman for technical assistance.

