## Supplementary Table 1 for "High throughput Characterization of *KCNB1* variants Associated with Developmental and Epileptic Encephalopathy"

**Supplementary Table 1.** Number of cells and p-values for the variant channel peak current density in comparison to that of WT. Values that differ from their respective WT are shown in red.

| Variant | Homomeric expression Ipeak density | | Co-expression with WT Ipeak density | |
|---|---|---|---|---|
| | n | p-value | n | p-value |
| **S202F** | 45 | 0.1348 | 68 | 0.3931 |
| **T210K** | 24 | 0.0037 | 44 | 0.8377 |
| **T210M** | 43 | 0.0002 | 29 | 0.6279 |
| **Y274fsX** | 43 | <0.0001 | 82 | 0.4954 |
| **R306C** | 27 | 0.0003 | 52 | 0.0363 |
| **R312C** | 25 | 0.0041 | 35 | 0.9998 |
| **R312H** | 31 | 0.0123 | 91 | 0.0957 |
| **R325Q** | 18 | 0.3151 | 43 | 0.7268 |
| **R325W** | 22 | 0.0589 | 29 | 0.9901 |
| **E330D** | 31 | <0.0001 | 67 | 0.0704 |
| **W370R** | 28 | <0.0001 | 69 | <0.0001 |
| **V378A** | 34 | 0.0007 | 29 | 0.9901 |
| **V378L** | 45 | <0.0001 | 78 | 0.0123 |
| **G381R** | 38 | <0.0001 | 88 | 0.0562 |
| **P385T** | 26 | <0.0001 | 64 | <0.0001 |
| **K391N** | 47 | <0.0001 | 44 | 0.1580 |
| **F416L** | 27 | <0.0001 | 64 | <0.0001 |
| **S457R** | 37 | 0.9999 | 47 | 0.9998 |
| **Y533X** | 29 | 0.0451 | 83 | 0.9994 |
