## Supplementary Figure 1 for "High throughput Characterization of *KCNB1* variants Associated with Developmental and Epileptic Encephalopathy"

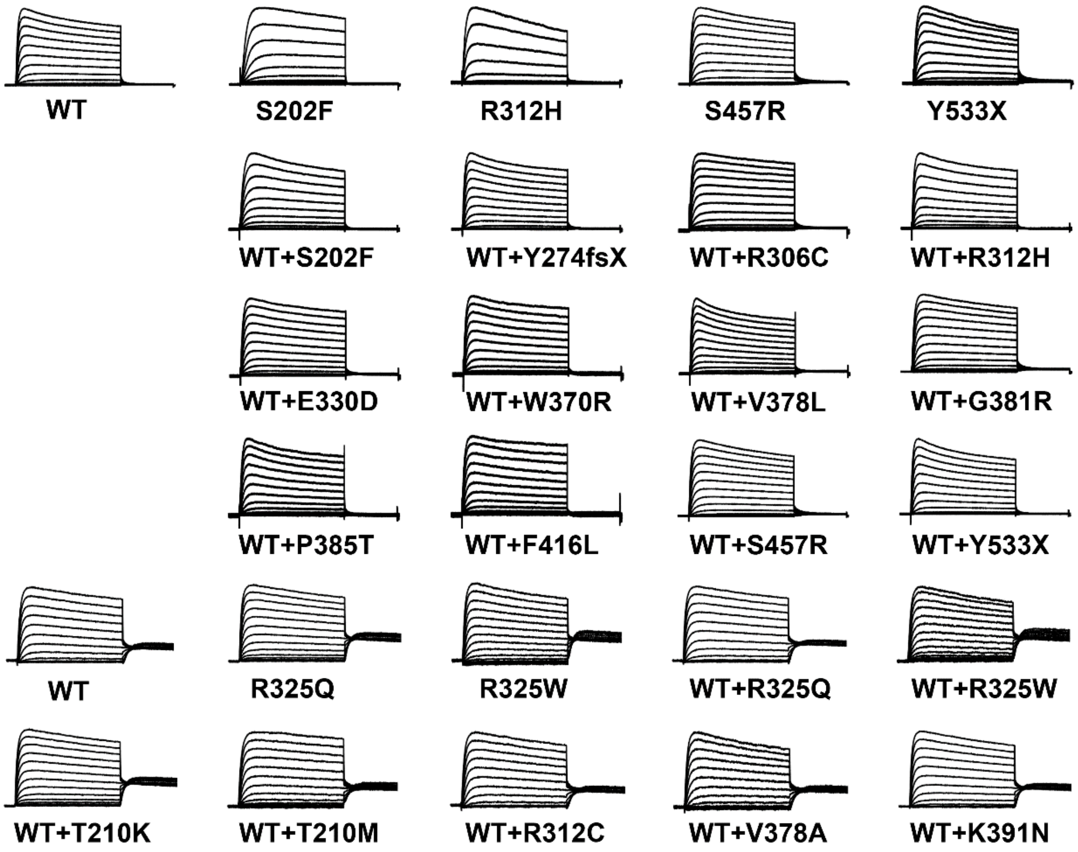


**Supplementary Figure 1.** Averaged current density traces normalized to the maximal peak current (at +60mV). For traces shown on the bottom two rows, the tail currents were held at 0 mV.
