## Supplementary Figure 2 for "High throughput Characterization of *KCNB1* variants Associated with Developmental and Epileptic Encephalopathy"

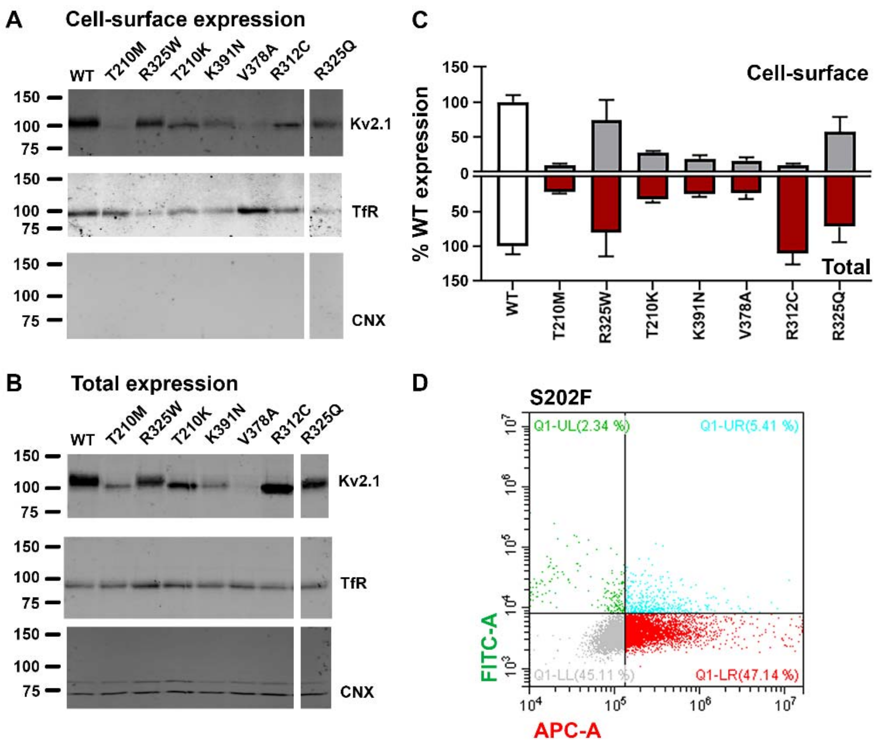


**Supplementary Figure 2.** Protein expression analyses of Kv2.1 variants. **A-B.** Western blot analyses of select Kv2.1 variants following cell-surface biotinylation. **C.** Quantification of western blot analyses shows reduced total expression level of Kv2.1 in variants; bands were normalized to transferrin receptor (TfR; loading control) and then compared to that of WT. **D.** The S202F variant shows protein expression yet reduced cell-surface expression possibly due to the steric hindrance between the epitope and the substituted phenylalanine.
